# PEACE: Prototype-aware Effector Analysis via Contrastive Embeddings

**DOI:** 10.64898/2026.04.19.719514

**Authors:** Xin Dai, Yuewei Lin, Shinjae Yoo, Qun Liu

## Abstract

Pathogenic fungi and oomycetes secrete effector proteins that manipulate host defenses and physiology to facilitate infection. However, in a typical secretome, effectors represent only a small fraction of proteins, creating extreme class imbalance and making deep-learning-based effector prediction prone to false positives. Here, we introduce PEACE (Prototype-aware Effector Analysis via Contrastive Embeddings), a lightweight pipeline that integrates ProtTrans (ProtT5) sequence embeddings with prototype-aware contrastive training to enhance effector identification. We benchmark PEACE against EffectorP 3.0 on two datasets with realistic class ratios: a fungal-only dataset and a combined fungi+oomycete dataset. PEACE outperforms EffectorP 3.0 on both datasets, while maintaining highly competitive recall. Post-hoc analysis shows that PEACE forms compact effector clusters against a well-dispersed non-effector background, which improves precision in the high-recall regime. These findings demonstrate that prototype-aware objectives, combined with curated data, can improve effector discovery for high-throughput screening in plant pathology and biotechnology.

## 1 Introduction

Fungal and oomycete effectors are secreted proteins that interact with host receptors and remodel host physiology, suppress immunity, and reshape host-associated microbiomes, thereby determining disease outcomes [1]. Effector proteins are often grouped by where they act: *apoplastic* effectors function outside of host cells, whereas *cytoplasmic* effectors enter host cells and target intracellular pathways and signaling hubs. In response to pathogens, hosts have evolved immune recetpors to against pathogenic effectors. The evolution of an effector’s sequence and function is rapid and dynamic, determining if a disease will occur. However, identifying them from all secreted proteins (secretome) remains a challenge and a high-priority task [2].

Several computational tools have been developed to predict effectors from secreted protein repertoires. EffectorP 3.0 [3] is widely used for fungal and oomycete effectors. It trains multiple Naïve Bayes and decision tree models on curated positives and secretome-derived negatives and aggregates them by soft voting to assign apoplastic, cytoplasmic, or non-effector labels. WideEffHunter [4] is a modular, pattern-driven pipeline intended to predict canonical and non-canonical effectors. It integrates three discrete predictive signals: (i) presence of curated effector-related motifs, (ii) presence of curated effector-related domains, and (iii) homology to a validated effector. More recently, machine-learning models that explicitly leverage pretrained protein language models (pLMs) have emerged. *POOE* [5] (oomycetes) couples ProtTrans [6] embeddings with a support vector machine (SVM) to predict oomycete effectors. *Fungtion* [7] builds on ESM-1b [8] embeddings to predict fungal effectors and perform downstream relationship analyses.

A central difficulty in effector analysis using machine-learning models is that effectors are rare among hundreds to thousands of secreted proteins in a secretome. Most training and benchmarking protocols rely on balanced or near-balanced datasets (e.g., effector vs non-effector of 1:1 to 1:3), which may overestimate performance and lead to excessive false positives. EffectorP 3.0 explicitly subsamples negatives during training, and predictions can be sensitive to secretion filtering and amplify false positives when applied to secretome data [3]. Likewise, pLM-based approaches, such as POOE and Fungtion, demonstrate strong accuracy in curated settings, but do not directly address the severely imbalanced data encountered in practice.

Contrastive learning offers a complementary strategy because it optimizes the geometry of the representation space, not only the final decision boundary. At a high level, contrastive objectives pull together multiple views of the same input while pushing apart representations of different inputs [9, 10]. In the present setting, this is attractive because the rare effector class should form a compact, separable minority, whereas the abundant non-effector class should remain broadly distributed to reflect its biological diversity.

In this work, we describe the PEACE workflow, which pairs ProtTrans embeddings with a lightweight Multi-Layer Perceptron (MLP) [11] and a *prototype-aware contrastive* objective tailored to the binary effector–non-effector setting. We benchmark PEACE on the *Fungtion* dataset [7] and on a newly curated Fungi+Oomycete dataset with realistic class ratios, and we analyze the learned representation geometry to explain why prototype-aware training improves performance under severe imbalance.

## 2 Results

### 2.1 PEACE workflow and prototype-aware contrastive learning

The PEACE workflow (Fig. 1) addresses the extreme class imbalance inherent in effector prediction by integrating protein language model embeddings with a prototype-aware learning objective. We utilized ProtTrans to generate embeddings for protein sequences, creating multiple “views” of each protein via stochastic dropout (Sec. 4.2).

**Figure 1:**
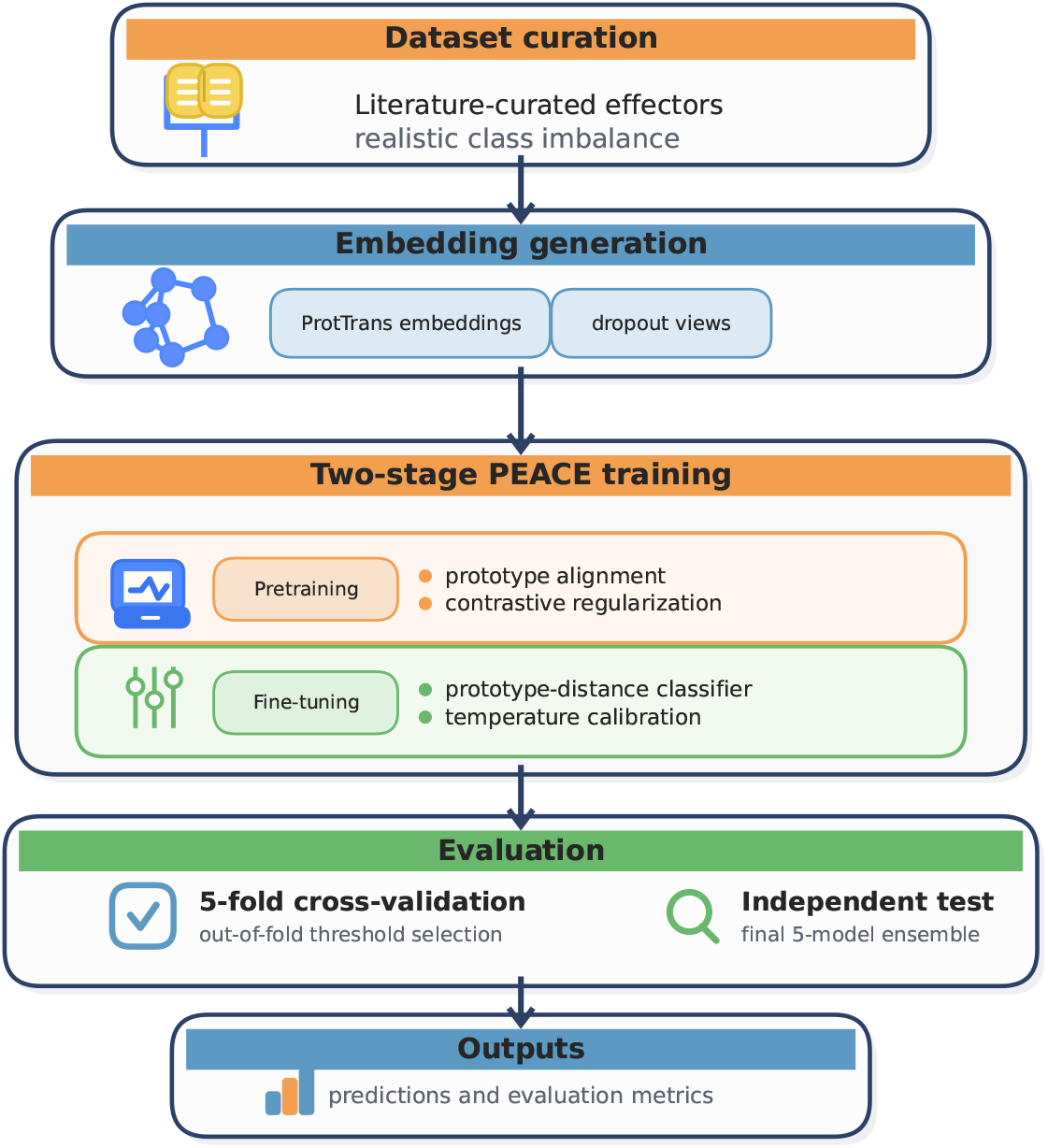
Overview of the PEACE workflow. Literature-curated datasets are embedded with Prot-Trans, followed by two-stage model training (pretraining with prototype alignment and contrastive regularization; fine-tuning with calibrated prototype-distance classification). The model performance evaluation includes a 5-fold cross-validation on the training set and an independent test set. Outputs include predictions and evaluation metrics.

To structure the latent space effectively, we adapted a supervised contrastive framework tailored for binary imbalance [12]. As illustrated in Fig. 2, the embedding space is anchored by two antipodal prototypes: **p**_1_ (effector) and **p**_0_ (non-effector). A key innovation is the use of *relative* similarity margins during training (Eq. (3)), which aggressively pulls minority-class effectors toward **p**_1_ while allowing the majority non-effectors to remain more dispersed. This geometry is optimized via a two-stage process: pretraining to align representations and fine-tuning to optimize a calibrated prototype-distance classifier.

**Figure 2:**
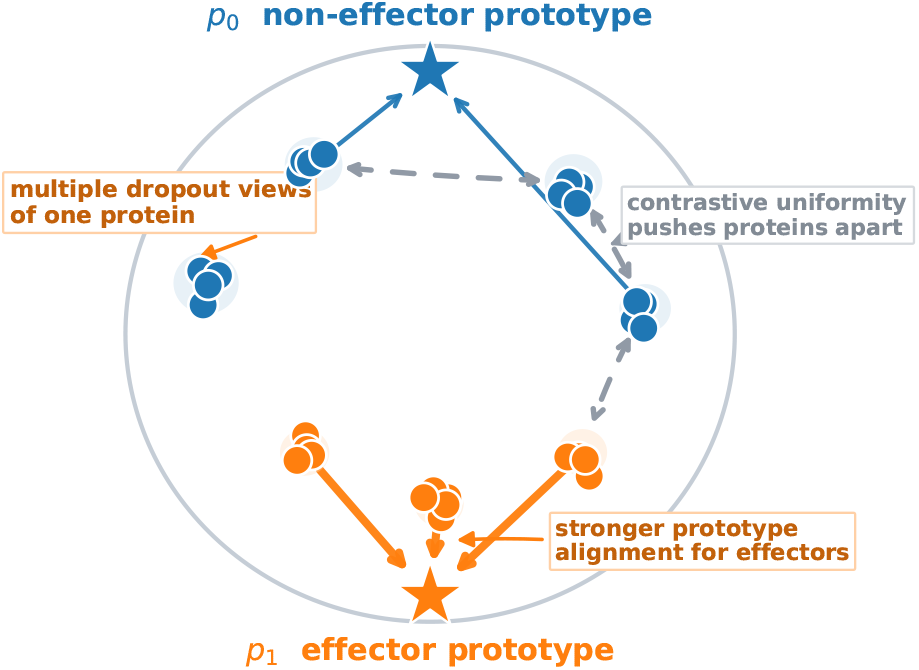
Schematic illustration of the PEACE prototype-aware contrastive learning framework in the embedding space (gray circle). The space is anchored by two class prototypes: non-effector *p*_0_ (blue star) and effector *p*_1_ (orange star). Each protein is represented by multiple stochastic dropout views, shown as a small cluster of circles of the same color. Prototype alignment pulls embeddings toward their corresponding class prototype (solid arrows), with stronger alignment for the minority effector class. At the same time, the contrastive objective promotes uniformity by pushing representations of different proteins apart (gray dashed arrows), improving class separation in the learned embedding space.

### 2.2 Benchmarking against state-of-the-art tools

We evaluated PEACE against widely used effector prediction tools on two independent held-out test sets representing realistic class ratios. We compared our performance to EffectorP 3.0 [3], the current standard for fungal/oomycete effectors, and WideEffHunter [4], a modular retrieval pipeline.

#### 2.2.1 Performance on fungal effectors (Fungtion dataset)

On the fungal-only Fungtion benchmark, PEACE substantially outperformed existing tools in handling realistic class imbalance (Table 1a). EffectorP 3.0, while achieving high recall (0.853), suffered from low precision (0.041), resulting in a low AUPRC of 0.142. This confirms that tools trained on balanced data often generate excessive false positives when applied to skewed secretomes. WideEffHunter similarly struggled with precision (0.010), rendering it unsuitable for genome-scale screening.

**Table 1:** Independent test performance comparing PEACE with external tools under realistic class imbalance. WideEffHunter outputs binary predictions (no calibrated scores), so ROC-AUC and AUPRC are not available. EffectorP 3.0 multiclass outputs are mapped to a binary effector label by treating *apoplastic* or *cytoplasmic* as effector.

| (a) Fungtion (fungal-only) |  |  |  |  |  |  |
| --- | --- | --- | --- | --- | --- | --- |
| Model | ROC-AUC | AUPRC | Prec. | Rec. | F1 | MCC |
| PEACE (Two-stage) | **0.963** | **0.558** | **0.217** | 0.676 | **0.329** | **0.372** |
| WideEffHunter | N/A | N/A | 0.010 | 0.706 | 0.019 | -0.026 |
| EffectorP 3.0 | 0.864 | 0.142 | 0.041 | **0.853** | 0.079 | 0.158 |

Table 1: Independent test performance comparing PEACE with external tools under realistic class imbalance.
| (b) Fungi+Oomycete |  |  |  |  |  |  |
| --- | --- | --- | --- | --- | --- | --- |
| Model | ROC-AUC | AUPRC | Prec. | Rec. | F1 | MCC |
| PEACE | **0.879** | **0.254** | **0.112** | 0.681 | **0.192** | **0.245** |
| WideEffHunter | N/A | N/A | 0.025 | **0.877** | 0.048 | 0.054 |
| EffectorP 3.0 | 0.732 | 0.067 | 0.037 | 0.754 | 0.070 | 0.101 |

In contrast, PEACE achieved an AUPRC of **0.558**, nearly a four-fold improvement over EffectorP 3.0, and the highest F1 score (0.329) and MCC (0.372). Figure 3(a–b) presents the precision-recall curves for the independent test set. In the high-recall regime (*R* ≥ 0.7), PEACE maintains significantly higher precision than EffectorP 3.0, demonstrating its utility for prioritizing high-confidence candidates in large-scale screens.

**Figure 3:**
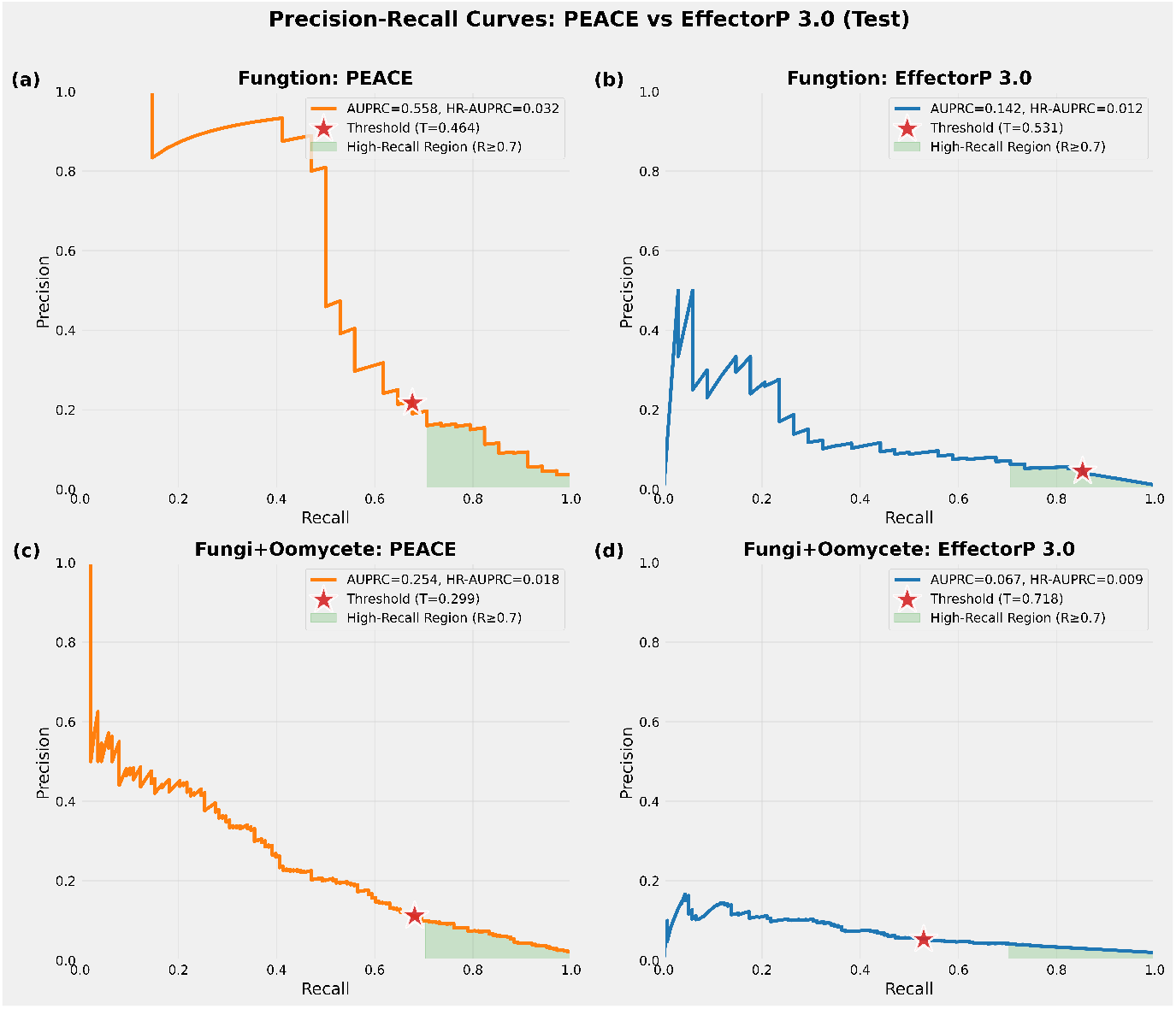
Precision-recall curves on independent test sets comparing PEACE and EffectorP 3.0. Top row (a–b): *Fungtion* dataset (fungal-only). Bottom row (c–d): Combined *Fungi+Oomycete* dataset. Light-green shading denotes the high-recall region (*R* ≥ 0.7). PEACE consistently maintains higher precision in the high-recall regime compared to EffectorP 3.0. The red star indicates the operating point derived from the global decision threshold selected by Youden’s *J* on validation data.

#### 2.2.2 Performance on combined Fungi+Oomycete effectors

We further evaluated the models on a newly curated, challenging Fungi+Oomycete dataset (Table 1b). This dataset introduces greater sequence diversity and distinct negative backgrounds (Sec. 4.1). Consistent with the fungal-only results, PEACE provided the best balance of performance metrics, achieving an AUPRC of **0.254** and an MCC of **0.245**. EffectorP 3.0 showed a marked performance drop on this combined set, with an AUPRC of 0.067 and MCC of 0.101. While WideEffHunter achieved the highest raw recall (0.877), its inability to control false positives (F1 0.048) limits its practical application.

Although the absolute performance of PEACE on the combined dataset was lower than on the fungal-only set (AUPRC 0.254 vs 0.558), it remained the only method to offer a viable trade-off between sensitivity and specificity (Fig. 3c–d). These results suggest that while taxon-specific models may be optimal, prototype-aware training provides robust generalization even in heterogeneous, highly imbalanced contexts.

### 2.3 Ablation studies

To validate the architectural and training choices in PEACE, we performed ablation studies comparing our proposed two-stage framework against two variants:

1. **Single-stage:** The same prototype-aware objective trained end-to-end without separate pretraining. See Eq. (5) in Sec. 4.4 for details.
2. **BCE Baseline:** A standard two-layer MLP trained with Binary Cross-Entropy on deterministic embeddings (no prototype alignment and contrastive views).

#### Cross-validation analysis

We first assessed model stability using stratified 5-fold cross-validation (Table 2). On both datasets, the prototype-aware variants (Two-stage and Single-stage) consistently outperformed the BCE baseline. On the Fungtion dataset, the Two-stage model achieved an AUPRC of 0.552*±*0.030 compared to 0.449*±*0.046 for the baseline. Similarly, on the Fungi+Oomycete dataset, the Two-stage model led with an AUPRC of 0.338 *±* 0.037 versus 0.309 *±* 0.056 for the baseline. While the Single-stage model was competitive, the Two-stage approach provided the most stable gains in AUPRC and MCC. By isolating the contrastive representation learning phase from the classifier calibration phase, the two-stage approach prevents the dominant negative class from prematurely biasing the decision boundary before a robust, prototype-anchored geometry is fully established.

**Table 2:** Ablation studies: 5-fold cross-validation performance. Mean *±* sd across folds.

| (a) Fungtion (fungal-only) |  |  |  |  |  |  |
| --- | --- | --- | --- | --- | --- | --- |
| Model | ROC-AUC | AUPRC | F1 | Prec. | Rec. | MCC |
| PEACE (Two-stage) | $0.923 \pm 0.016$ | **$0.552 \pm 0.030$** | $0.280 \pm 0.097$ | $0.176 \pm 0.080$ | $0.870 \pm 0.081$ | $0.346 \pm 0.076$ |
| PEACE (Single-stage) | **$0.929 \pm 0.021$** | $0.530 \pm 0.077$ | **$0.291 \pm 0.057$** | **$0.179 \pm 0.043$** | $0.844 \pm 0.085$ | **$0.354 \pm 0.044$** |
| BCE Baseline | $0.907 \pm 0.021$ | $0.449 \pm 0.046$ | $0.194 \pm 0.056$ | $0.111 \pm 0.037$ | **$0.894 \pm 0.062$** | $0.266 \pm 0.053$ |

| (b) Fungi+Oomycete |  |  |  |  |  |  |
| --- | --- | --- | --- | --- | --- | --- |
| Model | ROC-AUC | AUPRC | F1 | Prec. | Rec. | MCC |
| PEACE (Two-stage) | **$0.902 \pm 0.011$** | **$0.338 \pm 0.037$** | $0.188 \pm 0.023$ | $0.106 \pm 0.015$ | $0.827 \pm 0.036$ | **$0.264 \pm 0.020$** |
| PEACE (Single-stage) | $0.896 \pm 0.026$ | $0.312 \pm 0.057$ | **$0.193 \pm 0.022$** | **$0.110 \pm 0.014$** | $0.779 \pm 0.063$ | $0.262 \pm 0.028$ |
| BCE Baseline | $0.898 \pm 0.025$ | $0.309 \pm 0.056$ | $0.167 \pm 0.047$ | $0.094 \pm 0.031$ | **$0.830 \pm 0.055$** | $0.241 \pm 0.043$ |

#### Generalization to held-out test sets

The advantages of the Two-stage approach were further confirmed on the independent test sets (Table 3). On the Fungtion test set, Two-stage training yielded an AUPRC of 0.558, surpassing both the Single-stage (0.514) and the BCE baseline (0.540). The performance gap was clearer in the F1 and MCC metrics, where the Two-stage model (F1 0.329) significantly outperformed the baseline (F1 0.138). This trend held for the combined dataset, where the Two-stage model achieved the highest AUPRC (0.254) and MCC (0.245). These ablations demonstrate that while pLM embeddings are powerful on their own (as seen in the strong baseline performance relative to EffectorP), the specific combination of prototype alignment and staged optimization is critical for maximizing precision under severe imbalance.

**Table 3:** Ablation studies: Independent test set performance.

| (a) Fungtion (fungal-only) |  |  |  |  |  |  |
| --- | --- | --- | --- | --- | --- | --- |
| Model | ROC-AUC | AUPRC | Prec. | Rec. | F1 | MCC |
| PEACE (Two-stage) | **0.963** | **0.558** | **0.217** | 0.676 | **0.329** | **0.372** |
| PEACE (Single-stage) | 0.947 | 0.514 | 0.136 | 0.735 | 0.229 | 0.301 |
| BCE Baseline | 0.946 | 0.540 | 0.075 | 0.794 | 0.138 | 0.224 |

| (b) Fungi+Oomycete |  |  |  |  |  |  |
| --- | --- | --- | --- | --- | --- | --- |
| Model | ROC-AUC | AUPRC | Prec. | Rec. | F1 | MCC |
| PEACE (Two-stage) | **0.879** | **0.254** | **0.112** | 0.681 | **0.192** | **0.245** |
| PEACE (Single-stage) | 0.878 | 0.247 | 0.109 | 0.681 | 0.187 | 0.240 |
| BCE Baseline | 0.877 | 0.245 | 0.074 | 0.797 | 0.135 | 0.202 |

### 2.4 Representation geometry and the role of prototypes

To evaluate the role of prototype-aware contrastive learning in performance enhancement, we analyzed the embedding geometry on the *Fungtion* test set (Fig. 4, see Fig. S2 for the *Fungi+Oomycete* set).

**Figure 4:**
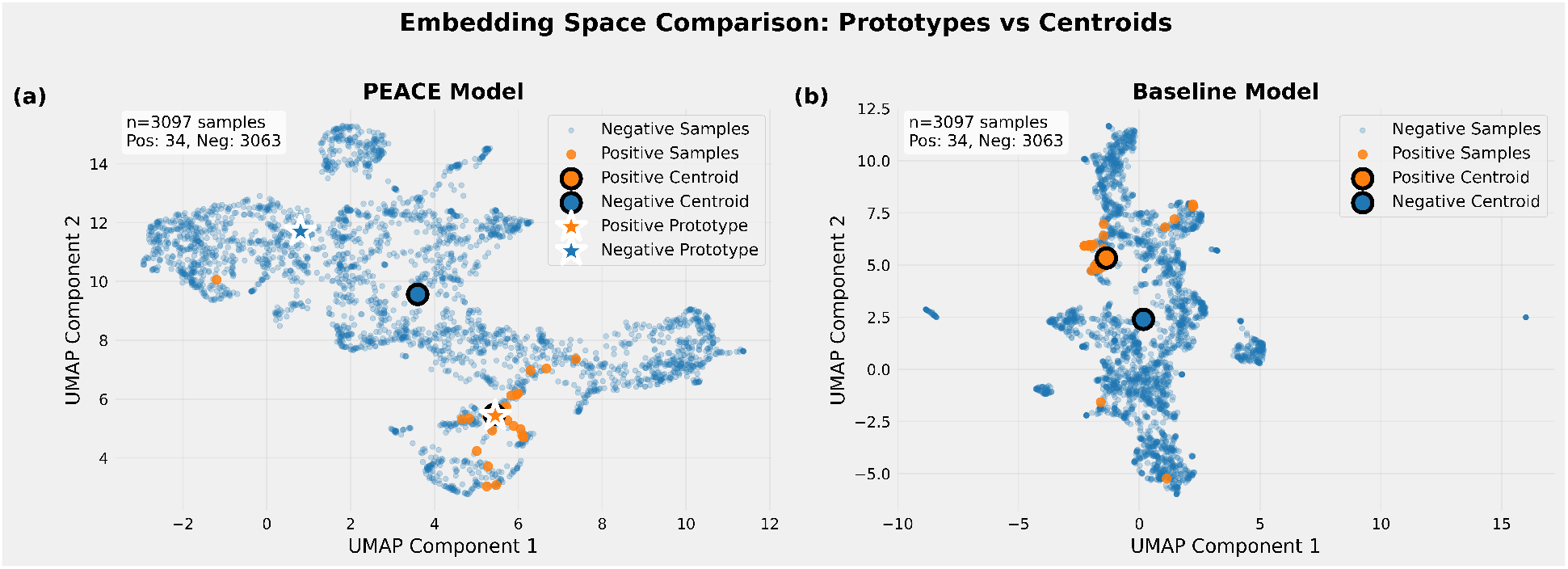
UMAP [14] visualization of learned embeddings on the *Fungtion* test set. Left: PEACE (Two-stage) model. The positive centroid (blue) aligns with the effector prototype **p**_1_, while negatives are dispersed, preventing mode collapse. Right: BCE Baseline model. Negatives collapse into a tight cluster, while positives are diffuse, leading to poorer discrimination.

#### Geometry anchored by prototypes

In our model, the positive class centroid aligns closely with the effector prototype (**p**_1_). This confirms that the Exponential Moving Average (EMA) updates during pretraining successfully stabilized the minority class representation. Conversely, the negative centroid is displaced from its antipodal prototype (**p**_0_), occupying a broad, diffuse region. This dispersion is a desirable property; it indicates that the unsupervised contrastive loss successfully prevented the majority class from collapsing into a tight cluster, preserving the intrinsic diversity of non-effectors.

#### Embedding uniformity prevents collapse

In contrast, the BCE Baseline produced a geometry where negatives formed a tight, dense cluster (Fig. 4, right). While this might imply structure, in a highly imbalanced setting, such tightness often reflects “feature collapse,” where the model learns a shortcut to map all non-effectors to a single mode. This is quantified by the Gaussian Potential Uniformity (GPU) [13] metric (Fig. 5 lower panels, see Eq. (1) for definition and Fig. S3 for the Fungi+Oomycete dataset). The baseline exhibits high GPU (low uniformity) for the negative class. Our model yields a significantly lower GPU for negatives, reflecting a more uniform distribution. Additionally, our model achieved very low inter-class similarity compared to the baseline (Fig. 5 upper panels), providing a cleaner separation margin that supports the calibrated classification required for high-precision effector discovery.

**Figure 5:**
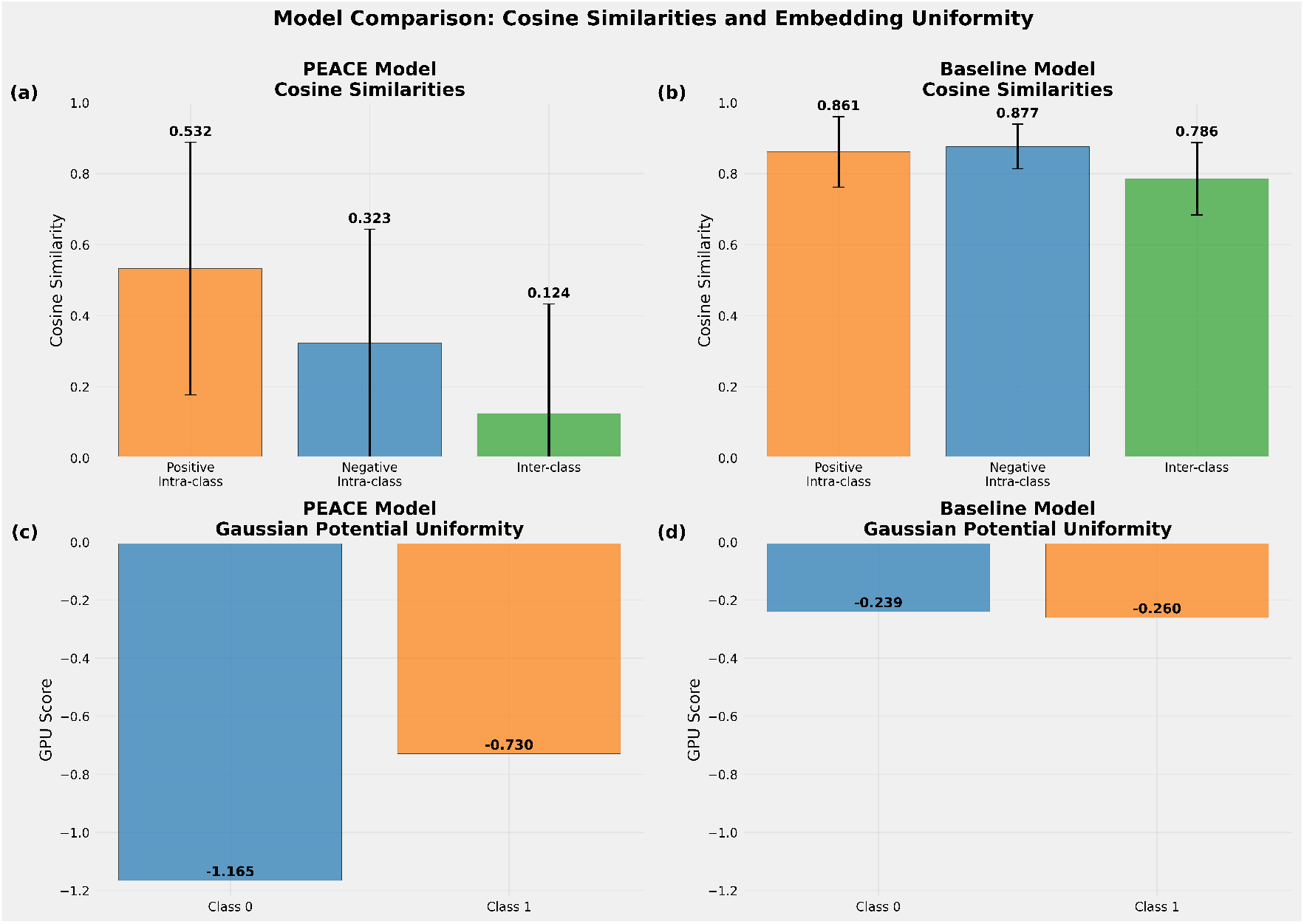
Comparison of embedding similarity statistics on the *Fungtion* test set. PEACE (top) achieves low inter-class similarity and high intra-positive similarity. The Baseline (bottom) exhibits uniformly high similarities across all partitions, reflecting class entanglement. Gaussian Potential Uniformity (GPU) scores indicate that PEACE maintains a more uniform (spread-out) distribution for the majority negative class compared to the baseline.

## 3 Discussion

### 3.1 Implications of testing on realistic class imbalance

Our benchmarking results highlight a critical disconnect in computational effector prediction: models optimized on balanced datasets often fail when deployed on genome-scale secretomes where effectors are rare [3, 5]. The stark difference in precision between PEACE and EffectorP 3.0 (Table 1) demonstrates that standard supervised learning often solves the “easy” problem of distinguishing effectors from random sequences, but fails the “hard” problem of distinguishing effectors from the vast, diverse background of non-effector secreted proteins.

The geometric analysis in Sec. 2.4 provides the mechanistic explanation for this improvement. By enforcing a prototype-anchored geometry, PEACE prevents the majority class from collapsing into a tight cluster—a common pathology in standard classifiers that we termed “feature collapse.” Instead, our approach forces the model to maintain a diffuse, uniform representation of non-effectors (Fig. 4). This suggests that for rare-event biological sequence prediction, the training objective must explicitly account for the *distributional asymmetry* of the classes, rather than simply re-weighting the loss or subsampling the data.

### 3.2 Generalization versus taxon-specificity

A notable finding was the performance gap between the fungal-only and the combined Fungi+Oomycete datasets. While PEACE outperformed baselines in both settings, its AUPRC was substantially higher on the fungal-only set (0.558) compared to the combined set (0.254). We attribute this to the biological heterogeneity of the targets; oomycete effectors often contain distinct translocation motifs (e.g., RxLR, CRN) that do not share semantic similarity with fungal effectors in the protein language model space.

This observation supports the “No Free Lunch” theorem in effector prediction: a single, omnibus model struggles to reconcile the distinct negative backgrounds and sequence motifs of distantly related taxa. The geometric entanglement is harder to resolve when the definition of “effector” spans multiple kingdoms. Consequently, we recommend that future pipelines adopt a hierarchical approach—first classifying the organism, then applying a taxon-specific effector predictor—rather than attempting to train a universal effector model.

### 3.3 Limitations: The dependency on upstream secretion filtering

A fundamental limitation of PEACE is the reliance on accurate upstream secretion prediction. Our datasets (and those of EffectorP) assume that the input proteins are correctly identified as secreted. However, SignalP [15] and similar tools have non-negligible error rates. If a non-secreted intracellular protein is erroneously passed to the effector predictor, it introduces a false positive source that current models are not trained to recognize.

The high-recall capability of existing tools often comes at the cost of amplifying these upstream errors [16]. Because PEACE is designed to maintain a high-precision threshold, it mitigates some of this risk, but it does not eliminate it. A distinct long-term goal for the community is the development of *joint* predictive frameworks that model secretion signals and effector properties simultaneously, thereby minimizing the propagation of errors from separate pipeline stages.

### 3.4 Future outlook: Integrating structure

While this work focused on maximizing the utility of sequence embeddings via better training objectives, the next frontier lies in multi-modal integration. Recent advances in structure prediction (e.g., AlphaFold2[17]/3[18], ESMFold [19]) offer orthogonal information to sequence data. Since effectors are often structurally conserved even when sequences diverge, structural embeddings could resolve cases where sequence homology is undetectable. Integrating structural modalities into the PEACE prototype-aware framework could further refine the embedding geometry, potentially closing the performance gap observed in the combined Fungi+Oomycete dataset.

## 4 Methods

### 4.1 Datasets and curation

Table 4 provides a high-level summary of the two datasets used in this study. **Fungtion dataset (fungal-only)**. We used the Fungtion benchmark from Li *et al*. [7] containing experimentally validated *fungal* positives aggregated from multiple sources (EffectorP, Effector–GAN, EffHunter) and negatives drawn from Effector–GAN [20]. Sequence identities were reduced at 40% using CD–HIT followed by structure-level filtering using ECOD [21], yielding compact training and test sets (Table 4b).

**Table 4:** Dataset statistics used in this study.

| (a) New Fungi+Oomycete dataset |  |  |  |  | (b) Fungtion dataset |  |  |  |  |
| --- | --- | --- | --- | --- | --- | --- | --- | --- | --- |
| Split | Total | Pos. | Neg. | Pos:Neg | Split | Total | Pos. | Neg. | Pos:Neg |
| Pretrain | 33,231 | 1,271 | 31,960 | 1:25 | Train | 6,918 | 178 | 6,740 | 1:38 |
| Train | 27,897 | 547 | 27,350 | 1:50 | Test | 3,097 | 34 | 3,063 | 1:90 |
| Test | 7,038 | 138 | 6,900 | 1:50 |  |  |  |  |  |

#### New Fungi+Oomycete dataset

To support model evaluation under realistic class imbalance and broader taxonomic coverage, we compiled a new *Fungi+Oomycete* dataset. Positives include (i) fungal effectors used in prior predictors and (ii) oomycete effectors from EffectorP 3.0 and POOE [3, 5]. Positives were clustered and filtered at 40% sequence identity. Negatives include from two curated sets: (i) EffectorP 3.0 secretomes depleted for particular effector classes (e.g., saprophytes, ectomycorrhizae, and RxLR/non-RxLR oomycetes), and (ii) Effector–GAN’s non-effectors drawn from secreted, non-pathogenic proteins [3, 20].

#### Partitioning and filtering pipeline

We first applied CD–HIT clustering separately to the positive and negative sets to reduce redundancy within the class, using a 40% identity threshold and minimum coverage constraints. To further minimize label noise, we rigorously filtered the negative background using BLASTP [22]; any negative sequence sharing ≥40% sequence identity and ≥60% alignment coverage with an *E*-value threshold of 1 × 10^−5^ against any positive sequence was discarded.

We then performed an 80/20 cluster-based split on the positive set—assigning entire clusters to either training or test—to prevent highly similar sequences from appearing in both. From each cluster, we selected the representative sequence to form the positives in the train and test partitions. To emulate real-world class imbalance, we augmented these sets with negative representatives to maintain a fixed 50:1 negative-to-positive ratio in both fine-tuning and test splits. These two sets comprise the final **train/test** dataset used for supervised learning and evaluation.

For **pretraining**, we expanded the positive set by including *all* sequences from the training clusters, allowing intra-cluster similarity, and applied BLASTP (using the same strict thresholds) to remove any sequence sharing homologous traits with any test positive. This ensured rigorous train–test data separation without homology leakage. Finally, we added remaining negatives not already used in the fine-tuning/test partitions to enrich the contrastive learning stage. By construction, the train partition is a strict subset of the pretrain partition. A flowchart of the construction of Fungi+Oomycete dataset is shown in Fig. S4.

### 4.2 Embeddings and Variant Generation

We addressed effector vs non-effector imbalance data by pairing ProtTrans embeddings with a two-layer MLP and a *prototype-aware contrastive* objective tailored to the binary effector–non effector setting. Inspired by, but distinct from, the supervised contrastive fixes of Mildenberger *et al*. [12] for imbalanced data, we (i) anchor the space with antipodal class prototypes, (ii) activate alignment based on a *relative* similarity margin with asymmetric tolerances that more strongly repel positives from the majority direction, (iii) update the minority (effector) prototype by EMA during pretraining to stabilize its geometry, and (iv) perform calibrated classification directly on temperature-scaled prototype-distance scores. These choices extend “Supervised Prototypes/Supervised Minority” ideas to protein embeddings and more realistic effector discovery, where maintaining majority-class uniformity while tightly aligning a sparse minority is essential.

We used ProtT5-XL-UniRef50 [6], a protein language model trained on approximately 50 million protein sequences, to derive fixed-length representations for input protein sequences. For each sequence, we extracted the hidden states from the last encoder layer, applied mean pooling across the sequence dimension, and then performed *ℓ*_2_ normalization. This yielded a single 1024-dimensional embedding vector per protein.

For contrastive learning, we adopt a SimCSE-style [10] variant generation scheme to create multiple stochastic views of each input sequence. Specifically, for each protein, we computed *V* = 8 embeddings: one deterministic embedding with dropout disabled and seven stochastic embeddings with dropout enabled at inference time. These embeddings serve as distinct views of the same underlying sequence and are treated as positive pairs in the contrastive objective. All stochasticity arises exclusively from the pretrained model’s native dropout (dropout rate = 0.1); no other data augmentation or input perturbation is applied. This strategy encourages the model to learn invariant features while enhancing robustness under class imbalance.

### 4.3 Evaluation Metrics

To assess predictive performance, we employed a range of classification evaluation metrics with particular attention to address the class imbalance issue.

#### F1 score

The F1 score is the harmonic mean of precision and recall:

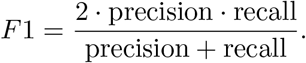

It balances recall to false positives and false negatives, but in highly imbalanced datasets it can still overemphasize precision.

#### Matthews Correlation Coefficient (MCC)

The MCC is defined as

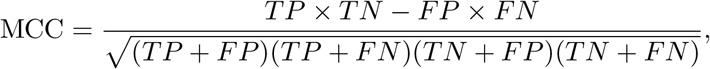

where *TP, TN, FP, FN* denote true positives, true negatives, false positives, and false negatives, respectively. MCC yields a single score in [−1, 1], with 1 indicating perfect prediction, 0 random performance, and −1 total disagreement. Because it incorporates all four entries of the confusion matrix, MCC is particularly informative even under strong class imbalance.

#### High-Recall AUPRC

The area under the precision–recall curve (AUPRC) is often more appropriate than ROC–AUC in rare-event settings. To focus on regions of practical interest, we computed partial AUPRC restricted to recall values above a threshold (e.g. *r* > 0.7). This emphasizes performance in the high-recall regime, where biological applications typically demand fewer false-positive effectors.

#### Youden’s *J* statistic

Threshold selection was guided by Youden’s *J* statistic, defined as

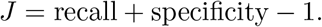

Although specificity tends to be trivially high in imbalanced problems, *J* provides a balanced criterion that avoids the severe recall penalties observed when optimizing directly for F1 or MCC.

Maximizing *J* typically shifts the decision boundary toward higher recall, thereby reducing false negatives at the cost of tolerating more false positives, which is a favorable trade-off in *in silico* effector screening.

#### Gaussian Potential Uniformity

Following Wang *et al*. [13], we quantified how evenly *ℓ*_2_-normalized embeddings *z*_*i*_ cover the hypersphere via the logarithm of the average pairwise Gaussian potential:

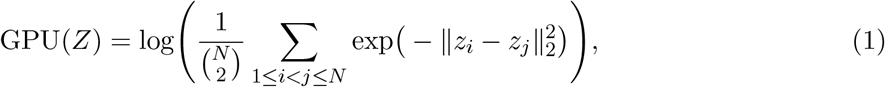

where lower values indicate greater uniformity (i.e., broader, less collapsed coverage). In imbalanced settings, a lower negative-class GPU reflects a more desirable, well-dispersed majority background.

### 4.4 Models and Loss Functions

#### Two-layer MLP architecture

Our model is a two-layer multilayer perceptron (MLP). The input is a fixed 1024-dimensional vector produced by ProtTrans embeddings. The first layer applies LayerNorm, a linear projection to 512 hidden units, GELU activation, and dropout (rate 0.2). The second layer again applies LayerNorm and linearly expands to 1024 dimensions. This 1024-dimensional output is *ℓ*_2_-normalized and serves as the contrastive embedding space used for prototype alignment and classification.

#### Baseline methods

For comparison, we implemented a similar two-layer MLP baseline: 1024→512 (LayerNorm + GELU + dropout) →1. This model produces a single scalar logit per protein, trained exclusively with binary cross-entropy. Unlike our prototype-based model, the baseline does not use a contrastive head, prototypes, or multiple variants; it relies solely on direct classification of single deterministic embeddings. At test time, predictions were obtained with a single forward pass and thresholded using the same calibration protocol.

#### Class prototypes and scoring

Two unit vectors, **p**_1_ (effector) and **p**_0_ (non-effector), anchor the embedding space. We initialized **p**_1_ as the centroid of positive-class embeddings and enforced antipodal symmetry by setting **p**_0_ = − **p**_1_. During both training and inference, each sequence *x* is represented by *V* stochastic views *z*^(1)^, …, *z*^(*V*)^. A sequence-level logit, Δ(*x*), is computed by averaging the temperature-scaled similarity differences across all *V* views. The final classification probability is obtained by applying the sigmoid function *σ* to this aggregated score:

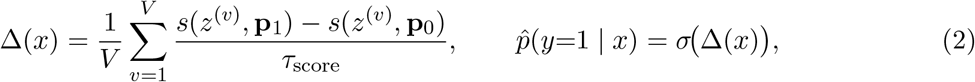

where *s*(·, ·) denotes cosine similarity.

#### Prototype alignment

Given a sequence with label *y* ∈ {0, 1} and a specific view *z*^(*v*)^, we denote the similarities to the prototypes as 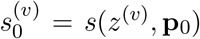 and 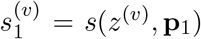. We use a two-class log-softmax loss to align the embeddings to their respective class prototypes:

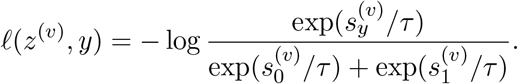

Alignment is *activated* only when the similarity margin over the opposite prototype is insufficient, utilizing asymmetric margins *ϵ*_pos_ > *ϵ*_neg_:

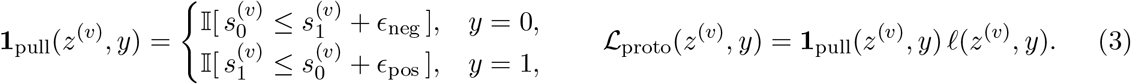

The total prototype alignment loss for a batch is aggregated by averaging over all views *z*^(*v*)^ that violate the margin constraints. Intuitively, the larger positive margin sustains a stronger attractive force that continues to pull minority-class (effector) embeddings toward the effector prototype until they are more tightly consolidated. In contrast, the smaller negative margin limits how aggressively majority-class samples are shaped, preventing the optimization from being dominated by abundant negatives.

#### Unsupervised contrastive regularizer

To maintain a global structure under class skew, we added an unsupervised InfoNCE contrastive loss [9] calculated over all *V* = 8 embedding variants of each sequence in a batch of size *N* :

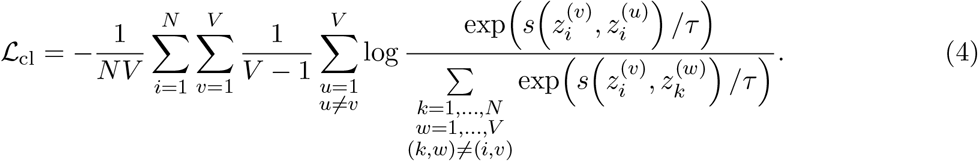

This loss encourages view-invariance and batch-wise uniformity, counteracts majority-class domination, and prevents collapse toward the majority prototype.

#### Fine-tuning objective

With prototypes frozen (or constrained), we optimized a temperature-scaled prototype-distance classifier using binary cross-entropy (BCE) applied to the aggregated sequence-level logit Δ(*x*), while retaining the stabilizing alignment and contrastive terms:

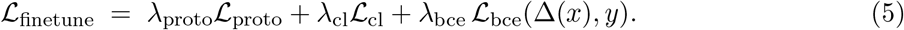

The individual loss weights *λ*_proto_, *λ*_cl_, and *λ*_bce_ serve as tunable hyperparameters.

### 4.5 Training Protocol and Hyperparameters

#### Two-stage training

Training proceeded in two stages: pretraining and fine-tuning. During the *pretraining* stage, representations were organized around the two prototypes using supervised alignment combined with an unsupervised contrastive term. To stabilize minority-class representation learning, the effector prototype (**p**_1_) was updated iteratively using an exponential moving average (EMA) of positive batch embeddings, while its non-effector counterpart remained antipodal. Pretraining ran for up to 25 epochs, with early stopping applied when the total loss failed to improve for 6 consecutive epochs. Learning rates followed a cosine annealing schedule with a small floor value to encourage smooth convergence.

In the *fine-tuning* stage, prototypes were frozen (or restricted by a warmup-freeze strategy) and the classifier was optimized directly on prototype distances with an added binary cross-entropy loss. This stage emphasizes calibrated decision boundaries while retaining prototype-informed structure. Fine-tuning ran for up to 100 epochs, with early stopping based on AUPRC on the validation set (patience of 8 epochs). A ReduceLROnPlateau scheduler adaptively decreased the learning rate when validation AUPRC plateaued, improving stability and preventing overfitting. Together, these staged objectives and adaptive schedules allowed the model to first learn robust, balanced representations before refining decision boundaries for classification.

#### Cross-validation and ensemble testing

We used stratified *k*-fold cross-validation to ensure balanced representations of positives across folds. For each fold, models were trained on *k* − 1 partitions and evaluated on the held-out validation fold. During training, decision boundaries were calibrated using Youden’s *J* statistic. To prevent data leakage and overfitting during threshold selection, out-of-fold (OOF) predictions were pooled across all *k* folds. A single global optimal threshold was then derived from this aggregated OOF distribution (see Fig. S1 for details). At test time, we ensembled the *k* = 5 trained models by averaging their probability outputs for each sequence, followed strictly by the application of the pre-calculated global threshold to obtain binary predictions.

#### Hyperparameter tuning

Key training parameters, including learning rates, weight decay, temperatures for contrastive and scoring objectives, prototype margins (*ϵ*_pos_, *ϵ*_neg_), and prototype momentum, were tuned using Bayesian optimization with a Tree-structured Parzen Estimator sampler. The optimization objective was the Matthews correlation coefficient (MCC) evaluated with global-thresholding, matching the final test protocol. Pretraining ran for up to 25 epochs, with early-stopping applied based on the total loss; while fine-tuning ran for 100 epochs, with early stopping applied based on validation MCC. Batch sizes of 1,024 sequences with 8 variants (one original plus seven dropout views) per sequence were used. The set of important hyperparameters can be found in Table S1.

## Data and Code Availability

Curated CSV files and training/evaluation code is available at https://github.com/Structurebiology-BNL/PEACE. For enhanced accessibility and user-friendly interaction, we have also implemented an easy-to-use Google Colab notebook, which serves as an inference server. It can be accessed at https://colab.research.google.com/github/Structurebiology-BNL/effector_prediction_with_contrastive_learning/blob/main/notebooks/fungus_inference_colab.ipynb.

## Acknowledgements

This work was supported by the U.S. Department of Energy (DOE), Office of Biological and Environmental Research (KP1601011).

## Notes

### Competing Interest Statement

The authors have declared no competing interest.

