## Supplemental Materials for "PEACE: Prototype-aware Effector Analysis via Contrastive Embeddings"

### Supplemental Materials For Prototype-Aware Contrastive Learning for Realistic Effector Screening

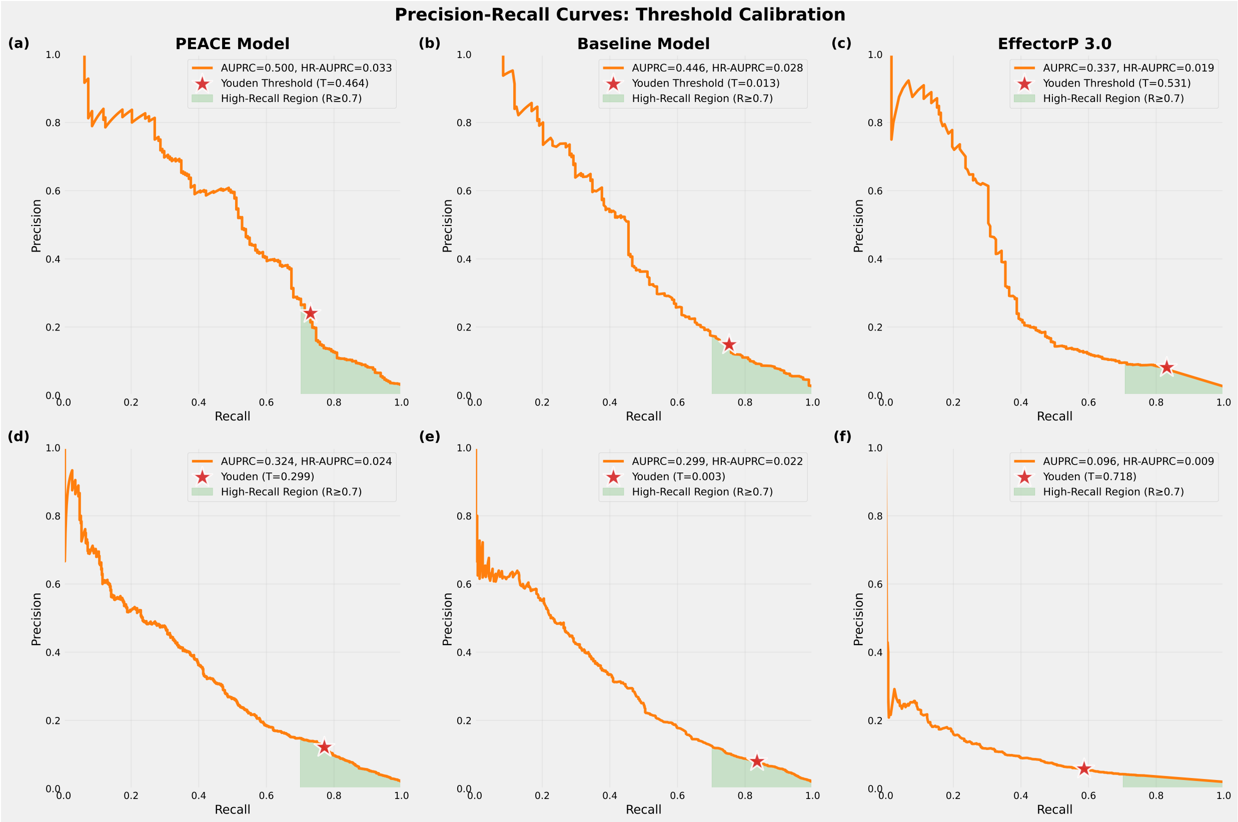


Figure 1: Precision–recall (PR) curves for decision threshold calibration. Top row (a–c): *Fungtion* dataset; bottom row (d–f): combined Fungi+Oomycete dataset. Columns (left to right) show our Prototype model (two-stage), the BCE Baseline, and EffectorP 3.0. For the Prototype and BCE Baseline models, curves are computed from pooled out-of-fold (OOF) predictions across 5 cross-validation folds. Because EffectorP 3.0 is a pre-trained external tool, its curves are generated by evaluating the tool directly on the full training set. Light-green shading denotes the high-recall region ($R \geq 0.7$). Legends report AUPRC, high-recall AUPRC (HR-AUPRC; Methods), and the decision threshold chosen by maximizing Youden's $J$; the red star marks this threshold on each curve. For our trained models, this global threshold derived from the OOF predictions is the one applied to the independent test sets in Table 1 and Fig. 3 of the main text.

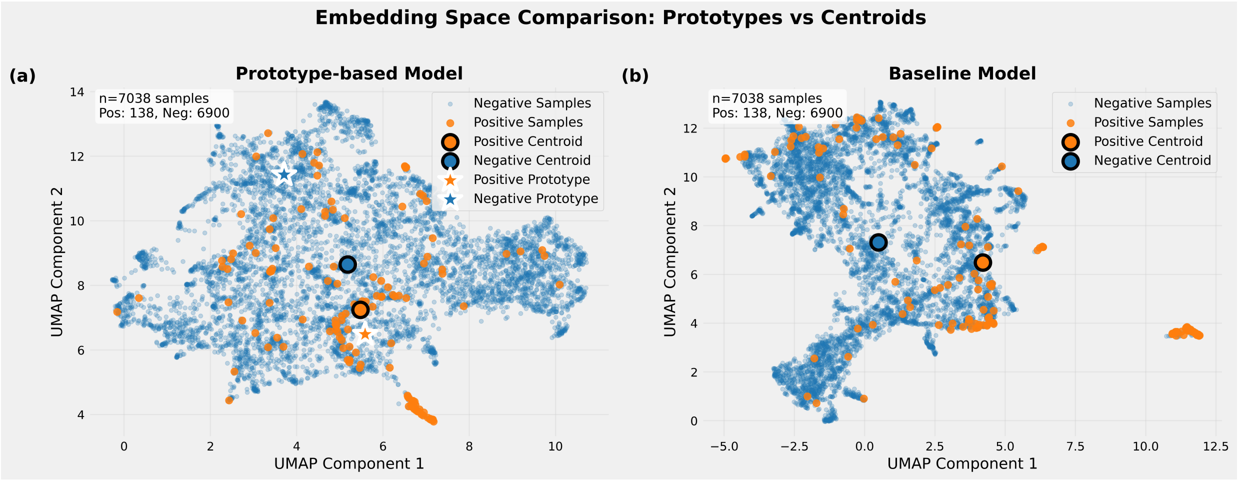


Figure 2: UMAP visualization of learned embeddings on the Fungi+Oomycete test set. Left: our model, with prototypes $(\mathbf{p}_1, \mathbf{p}_0)$ and class centroids marked. The positive centroid closely aligns with $\mathbf{p}_1$, while the negative centroid is displaced from $\mathbf{p}_0$, and negatives cover a broad region. Right: baseline model, where negatives collapse into a tighter cluster and positives are more diffuse.

| Setting | Effector dataset | Fungtion dataset |
|---|---|---|
| Batch size | 1024 | 1024 |
| Variants per sequence | 7 + deterministic | 7 + deterministic |
| Pretraining epochs | 25 | 25 |
| Finetuning epochs | 100 | 100 |
| $\tau$ | 0.071 | 0.052 |
| $\tau_{\text{score}}$ | 0.192 | 0.342 |
| $(\epsilon_{\text{pos}}, \epsilon_{\text{neg}})$ | $(0.235, 0.097)$ | $(0.215, 0.015)$ |
| Prototype update momentum | 0.97 | 0.95 |

Table 1: Representative tuned hyperparameters for the best models by dataset. $\tau$ = contrastive temperature; $\tau_{\text{score}}$ = scoring temperature for prototype-distance; $(\epsilon_{\text{pos}}, \epsilon_{\text{neg}})$ = relative-similarity margins that gate prototype alignment; "Prototype update momentum" = EMA coefficient used to update the positive prototype during pretraining.

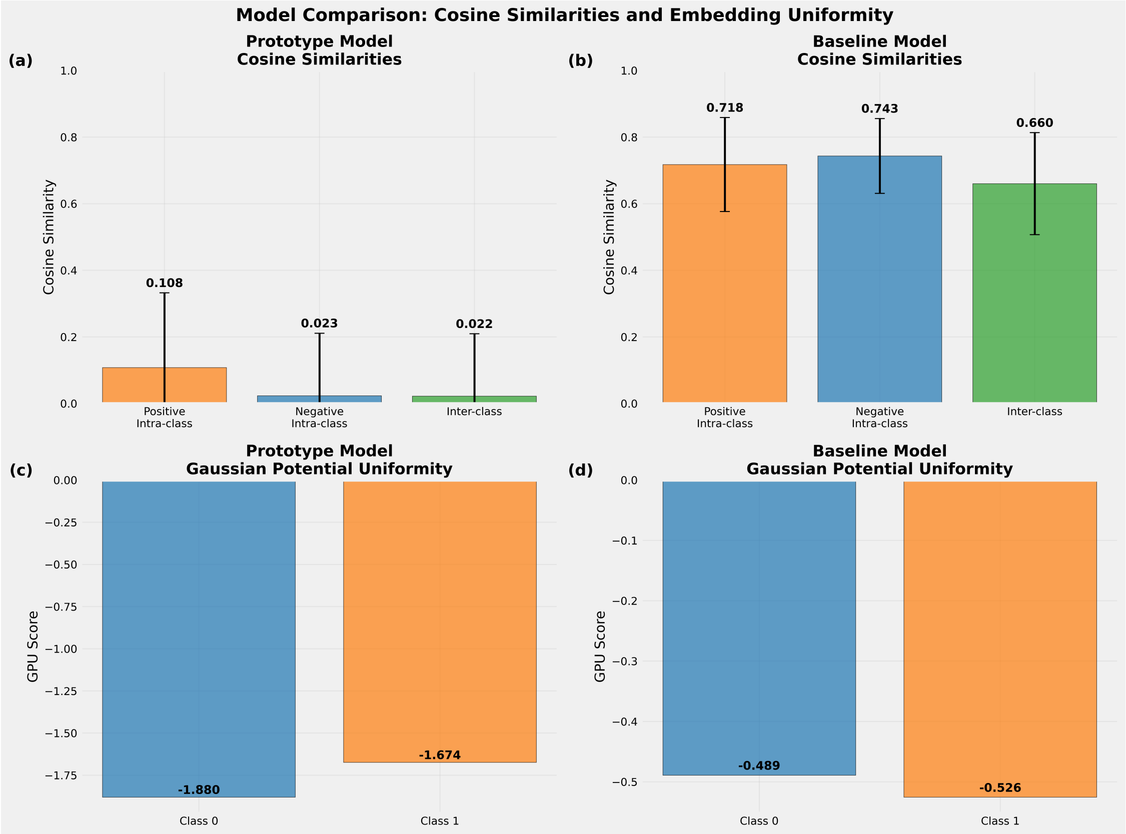


Figure 3: Comparison of embedding similarity statistics on the *Fungi+Oomycete* test set between our prototype-based model and the BCE baseline. Our model achieves higher intra-positive similarity than intra-negative similarity and very low inter-class similarity. The baseline exhibits uniformly high similarities, reflecting class entanglement. Gaussian Potential Uniformity (GPU), explained in the text, further demonstrates that negatives in our model are more uniformly distributed, preventing collapse and improving generalization.

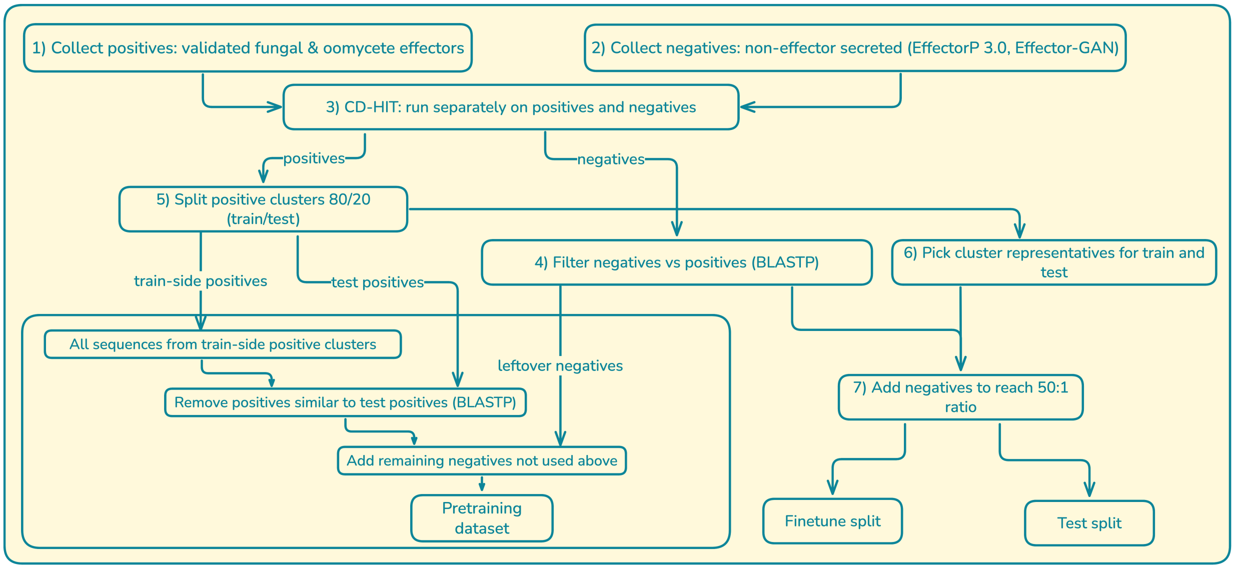


Figure 4: Overview of the dataset construction pipeline used for the new Fungi+Oomycete dataset.
